# Exosomally targeting microRNA23a ameliorates microvascular endothelial barrier dysfunction following rickettsial infection

**DOI:** 10.1101/2022.03.25.485825

**Authors:** Changcheng Zhou, Jiani Bei, Yuan Qiu, Qing Chang, Emmanuel Nyong, Jun Yang, Balaji Krishnan, Kamil Khanipov, Yang Jin, Xiang Fang, Angelo Gaitas, Bin Gong

## Abstract

Spotted fever group rickettsioses caused by *Rickettsia* (*R*) are devastating human infections, which mainly target microvascular endothelial cells (EC) and can induce lethal EC barrier dysfunction in the brain and lungs. Our previous evidence reveals that exosomes (Exos) derived from rickettsial-infected ECs, namely *R*-ECExos, can induce disruption of the tight junctional (TJ) protein ZO-1 and barrier dysfunction of human normal recipient brain microvascular endothelial cells (BMECs). However, the underlying mechanism remains elusive. Given that we have observed that microRNA23a (miR23a), a negative regulator of endothelial ZO-1 mRNA, is selectively sorted into *R*-ECExos, the aim of the present study was to characterize the potential functional role of exosomal miR23a delivered by *R*-ECExos in normal recipient BMECs. We demonstrated that EC-derived Exos (ECExos) have the capacity to deliver oligonucleotide RNAs to normal recipient BMECs in an RNase-abundant environment. miR23a in ECExos impairs normal recipient BMEC barrier function, directly targeting TJ protein ZO-1 mRNAs. In separate studies using a traditional *in vitro* model and a novel single living-cell biomechanical assay, our group demonstrated that miR23a anti-sense oligonucleotide-enriched ECExos ameliorate *R*-ECExo-provoked recipient BMEC dysfunction in association with stabilization of ZO-1 in a dose-dependent manner. These results suggest that Exo-based therapy could potentially prove to be a promising strategy to improve vascular barrier function during bacterial infection and concomitant inflammation.

## Introduction

Spotted fever rickettsioses are devastating human infections[1] that are caused by obligate intracellular bacteria of the genus *Rickettsia* (*R*). Microvascular endothelial cells (ECs) are the primary targets of infection, and edema resulting from EC barrier dysfunction occurs in the brain and lungs in most cases of lethal spotted fever group rickettsial infections in humans[2]. A licensed vaccine is not available. Typically, rickettsial infection is controlled by appropriate antibiotic therapy if diagnosed early[3; 4]. However, rickettsial infections can cause nonspecific signs and symptoms rendering early clinical diagnosis difficult[5; 6]. Untreated or misdiagnosed infections with rickettsiae are frequently associated with severe morbidity and mortality[4; 6]. Although doxycycline is the antibiotic of choice for rickettsial infections, it only stops the bacteria from reproducing, but does not kill the rickettsiae[7; 8]. A fatality rate as high as 32% has been reported in hospitalized patients with Mediterranean spotted fever[9]. It is forecasted that global climate change will lead to more widespread distribution of rickettsioses[10; 11]. Comprehensive understanding of rickettsial pathogenesis is urgently needed for the development of novel therapeutics.

Extracellular vesicles (EVs) transfer functional mediators to neighboring and distant recipient cells[12; 13; 14; 15]. EVs are broadly classified into two categories, small particles (also known as Exos) and large particles (or microvesicles), distinguished by their intra-cellular origin[16; 17]. Recent evidence shows that senescence predominantly induces alterations in Exos, not microvesicles[18]. The formation of Exos is initiated by budding into the late endosome; the multivesicular body transits towards and fuses with the plasma membrane before releasing intraluminal vesicles into the extracellular environment as Exos[16; 19]. Exos contain many types of biomolecules[16] and convey signals to a large repertoire of recipient cells either locally or remotely by ferrying functional cargos, thus contributing to disease pathogenesis[20; 21; 22; 23; 24; 25]. We recently unveiled a novel functional role played by EC-derived exosomes (ECExos) during rickettsial infection[26]. We validated that ECs efficiently take up exosomes (Exos) *in vivo* and *in vitro*[26]. We found that rickettsial-infected ECExos (*R*-ECExos) induced disruption of the tight junctional (TJ) protein ZO-1 and barrier dysfunction of human normal recipient brain microvascular endothelial cells (BMECs) in a dose-dependent manner. This effect is dependent on exosomal RNA cargos[26]. However, the underlying mechnism remains unclear.

The EC barrier properties are structurally determined by junctional complexes between ECs, mainly the adherens junctions (AJs) and TJs[27; 28]. *In vitro* data obtained by transfection with “naked” oligonucleotide-mimics (i.e., unmodified and lacking a carrier) in a ribonuclease (RNase)-free environment, which are not naturally adsorbed, demonstrated that microRNA23a (miR23a) is a negative regulator of EC ZO-1[29]. Although microRNA (miR) species (i.e., less than 25 nucleotides) are relatively stable when compared with other RNA molecules, they remain vulnerable to RNase-mediated digestion in the extracellular space[30; 31; 32]. The discovery of extracellular miRs in the blood, despite the abundant presence of RNases, led to the proposal of a scenario in which miRs are encapsulated in EVs [30; 33; 34] or form circulating ribonucleoproteins [35; 36]. A growing number of reports have established that many, if not all, of the effects of EVs are mediated by miR cargos[34; 37; 38; 39; 40; 41; 42; 43], which remain functional to regulate cellular behaviors of the recipient cells[44]. Exo-mediated functional transfer of miRs has been reported with a broad range of downstream effects on recipient cells[45]. We observed that miR23a is selectively sorted into *R*-ECExos[26]. However, we did not detect different levels of miR23a in parent cell samples following infections[26; 46].

The aim of the present study is to characterize the potential functional role of exosomal miR23a, which was identified as being predominantly expressed in *R*-ECExos, resulting in endothelial hyperpermeability. We found the delivery of miR23a in Exos was efficiently taken up by normal recipient ECs, impairing their barrier structures and leading to hyperpermeability. Exosomal miR23a targets ZO-1 mRNAs in recipient ECs. Furthermore, delivery of anti-sense oligonucleotides (ASOs) of miR23a in Exos to recipient cells during normal culture can ameliorate *R*-ECExo-induced endothelial barrier dysfunction, thus providing evidence for a potential Exo-based therapeutic to improve vascular barrier function during infection and concomitant inflammation.

## Results

### 1. ECExos can deliver miR23a mimic oligonucleotides to recipient BMECs in the presence of RNases

Size-exclusion chromatography (SEC) was utilized to isolate ECExos from human umbilical vein EC (HUVEC) culture media that had been passed through two 0.2 μm filters[26]. *R*-ECExos were purified 72 hrs post-infection from *R. parkeri*-infected HUVECs; a multiplicity of infection (MOI) of 10 was used. Quantitative real-time PCR validated that no rickettsial DNA copies were detected in the *R*-ECExo material (**Fig. 1A**). Sizes and morphologies of isolated particles were evaluated using atomic force microscopy (AFM)[26] and Tunable Resistive Pulse Sensing (TRPS) nanoparticle tracking analysis (NTA) (qNano Gold, Izon, Medford, MA)[47; 48] (**Fig. 1B, D**). The size distribution of isolated EVs was confirmed to be in the range of 50 nm to 150 nm, which is the expected size of Exos. We also verified the purity of isolated Exos using western immunoblotting to detect traditional exosomal markers as shown in **Fig. 1C**. As we reported, our purified ECExos and *R*-ECExos were free of bacteria or bacterial DNA, were intact, and did not aggregate.

**Figure 1.**
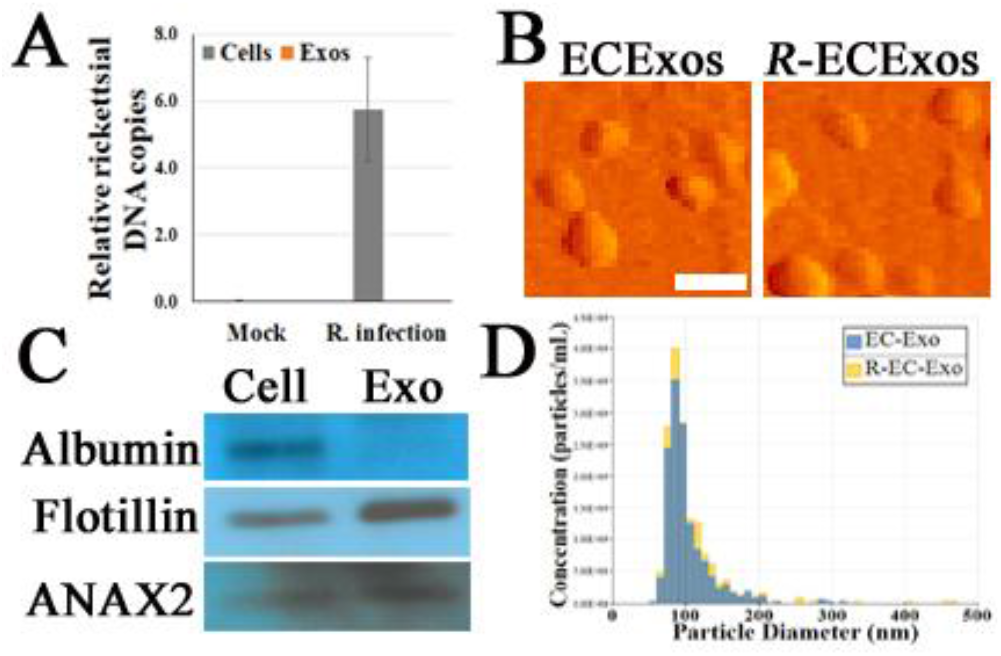
Characterization of HUVEC-derived ECExos using SEC isolation. **A**, Quantities of rickettsiae in parent ECs and ECExos (*n* = 3/group), determined by quantitative real-time PCR. Data are presented as means ± standard errors. **B**, ECExo morphology was verified using AFM (scale bars, 200 nm). **C**, Expression of three protein markers in 100 μg protein derived from ECExos was examined using western immunoblotting. **D**, Vesicle size distribution of isolated EVs was analyzed using NTA.

We reported that ECExos were absorbed by normal recipient BMECs [26]. To validate the capability of ECExos to deliver miRs to recipient cells, miR23a mimic oligonucleotides were conjugated with Cy3 fluorescent dyes[49], in which the fluorescent intensity is dependent on the nucleotide sequence[49]. Using a calcium-mediated miR-loading protocol that is specific for EVs[50], Cy3-conjugated miR23a mimic oligonucleotides were loaded into ECExos (2 pmol oligonucleotides/1 x 10^8^ Exos). Compared with the naked Cy3-conjugated group, the fluorescent signal was detected robustly in the ECExo group, suggesting that ECExos provide the miR23a with a shield against degradation by abundant RNases in normal EC culture media (**Fig. 2A**). A stem-loop PCR assay[26; 51; 52] showed statistically significant augmented levels of miR23a in recipient BMECs following exposure to miR23a mimic-enriched ECExos in normal culture media, compared with other groups (**Fig. 2B**).Therefore, ECExos have the capacity to deliver functional miR23a into normal recipient BMECs.

**Figure 2.**
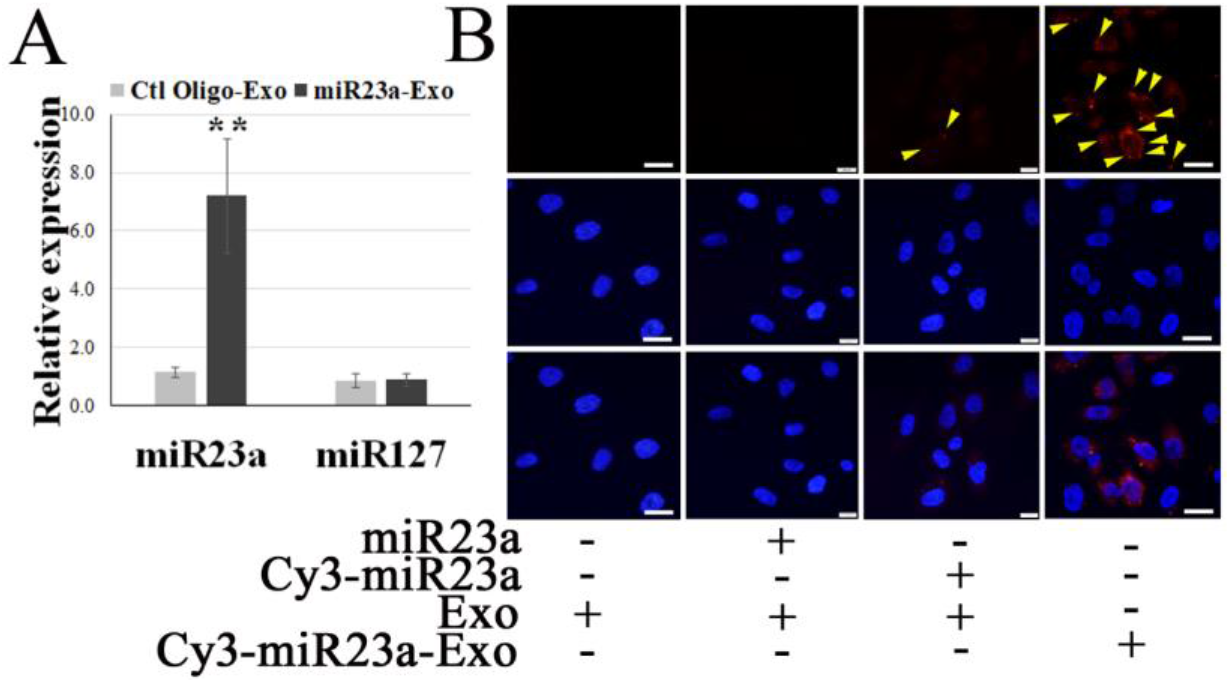
ECExos can deliver miR23a mimic oligonucleotides to recipient BMECs in the presence of RNases. **A**, Stem-loop PCR of miR23a and miR127 in miR23a-enriched ECExos (miR23a-Exo) or negative control oligonucleotides-enriched ECExos (Ctl Oligo-Exo) using a published protocol[50]. **, *p*<0.01. **B**, Fluorescent tracking of naked mir-23a or miR23a-Exo in BMECs in normal culture media after treatment for 12 hr.

### 2. Exosomal miR23a impairs the barrier function of recipient BMECs

Modulation of EC barrier function by miR23a has been documented[29; 53]. To evaluate the functional role that endothelial exosomal miR23a plays in recipient cells, the efficacy of miR23a mimic-enriched ECExos on the barrier function of recipient BMECs was examined. Compared with naked miR23a and negative control oligonucleotides (Ctl Oligo)-ECExos, we observed that miR23a mimic-enriched ECExos induced an increase in fluorescein isothiocyanate (FITC)-dextran[46] and a decrease in TEER[26] in recipient BMECs at 72 hrs post-exposure. (**Fig. 3**), suggesting that miR23a in ECExos impairs normal recipient BMEC barrier function.

**Figure 3.**
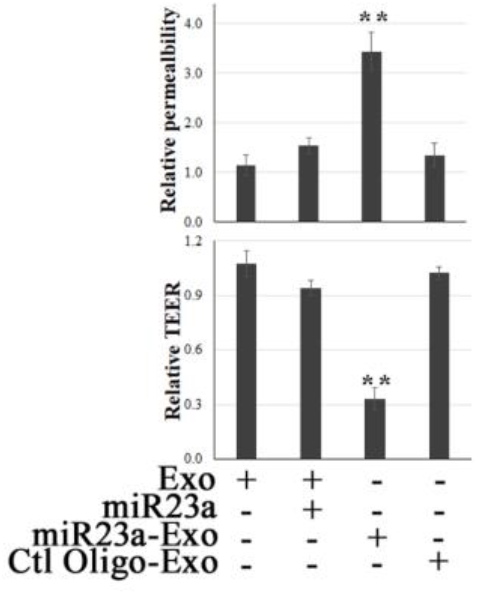
Exosomal miR23a impairs the barrier function of recipient BMECs. FITC-dextran-based assays[46] and TEER measurements[26] were performed in normal human BMECs following treatment with naked miR23a mimics (2 x 10^-5^ pmol/cell), ECExos (1,000 particles/cell), Ctl Oilgo-, or miR23a mimic-enriched ECExos (miR23a-Exo) (1,000 particles/cell) for 72 hrs. **, *p*<0.01.

### 3. Exosomal miR23a directly targets ZO-1 in recipient BMECs

It is reported that miR23a targets the TJ protein ZO-1 in experimentally transfected ECs[29; 53], although the question of natural adsorption of miR23a in RNase-environment had yet to be investigated. We examined ZO-1 mRNA levels and paracellular protein levels in recipient BMECs to determine the potential target of miR23a in the context of delivery by Exos in normal culture media.

Using a luciferase assay, we confirmed that ZO-1 is the direct molecular target of exosomal miR23a in recipient BMECs (**Fig. 4A**). We further observed that miR23a mimic-enriched ECExos downregulate ZO-1 mRNA levels (**Fig. 4B**) and disrupt paracellular ZO-1 in recipient BMECs 72 hr post-exposure (**Fig. 4C**), compared to Ctl Oligo-enriched ECExos. These data provide strong evidence that ZO-1 is the direct target of exosomal miR23a in recipient BMECs.

**Figure 4.**
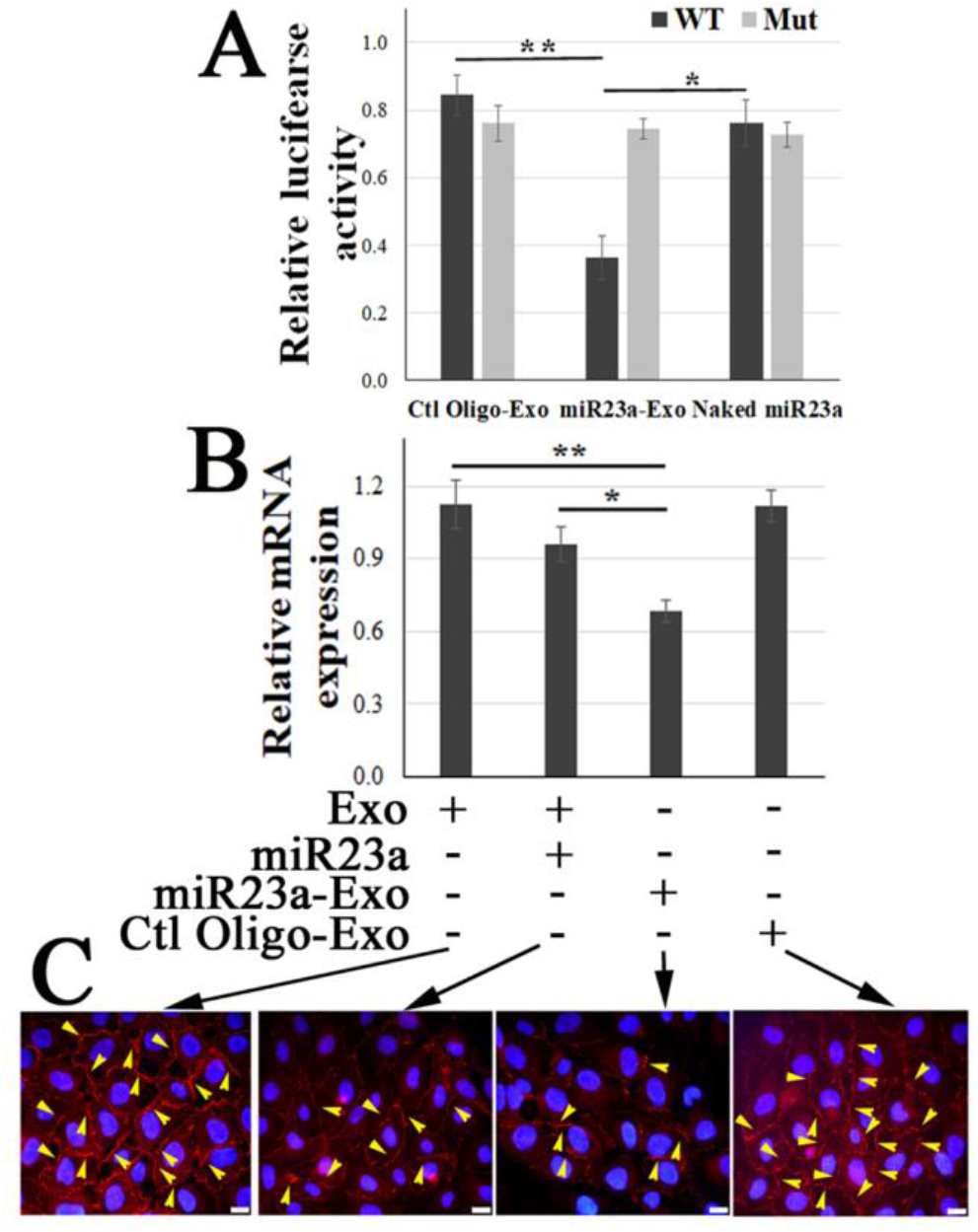
Exosomal miR23a directly targets ZO-1 of recipient BMECs. **A**, Luciferase reporter assay in which BMECs were exposed to naked miR23a mimics (2 x 10^-5^ pmol/cell), Ctl Oilgo-, or miR23a mimic-enriched ECExos (miR23a-Exo) (1,000 particles/cell) soon after transfection with reporter constructs containing full length 3’ UTR wild-type ZO-1 (WT) or mutated miR23a binding sites (Mut). Luciferase activity was quantified at 24 hrs post exposure. *N* = 3 independent experiments. *, *p*<0.05; **, *p*<0.01. **B**, Relative expression of ZO-1 mRNA by RT-PCR of recipient BMECs after exposure to naked miR23a mimic (2 x 10^-5^ pmol/cell), Ctl Oilgo-, or miR23a mimic-enriched ECExos (miR23a-Exo) (1,000 particles/cell) for 72 hr. **C**, ZO-1 immunofluorescence in recipient BMECs after exposure to naked miR23a mimics (2 x 10^-5^ pmol/cell), Ctl Oilgo-, or miR23a mimic-enriched ECExos (miR23a-Exo) (1,000 particles/cell) for 72 hr.

### 4. miR23a ASO-enriched ECExos ameliorate *R*-ECExo-provoked recipient BMEC dysfunction in association with stabilization of ZO-1

We reported endothelial barrier dysfunction and disruption of paracellular ZO-1 in recipient BMECs of *R*-ECExos following a detectable upregulation of miR23a in *R*-ECExos. Sequence-based ASOs have been shown to have pharmacological effects in the context of infection and inflammation in a manner dependent upon the target RNA[54; 55]. To evaluate the efficacy of miR23a ASO-enriched Exos on stabilizing barrier function of the recipient BMECs following exposure to *R*-ECExos, miR23a ASOs were loaded into normal ECExos derived from HUVECs using same protocol as for the mimic described above.

#### 4.1

We observed that miR23a ASO-enriched Exos (1,000 particles/cell) improved *R*-ECExo-induced recipient BMEC barrier dysfunction as evidenced by reducing FITC-dextran leakage (**Fig. 5A**) and increasing TEER measurements (**Fig. 5B**), both of which indicate improved cell barrier permeability and integrity.

**Figure 5.**
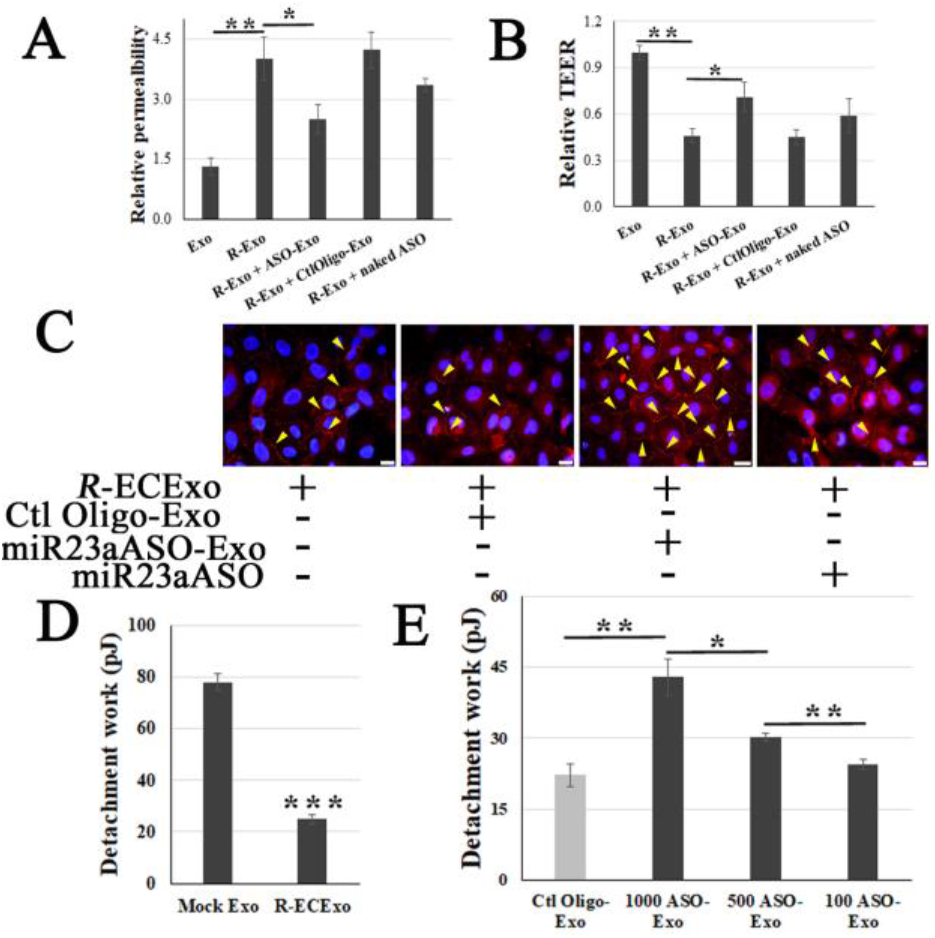
miR23a ASO-enriched ECExos ameliorate *R*-ECExo-provoked recipient BMEC dysfunction in association with stabilization of ZO-1. **A** and **B**, compared to naked miR23a ASO (naked ASO) and Ctl Oligo-enriched ECExos (Ctl Oligo-Exos), miR23a ASO-enriched ECExos (ASO-Exo) attenuate *R*-ECExo- (*R*-Exo-) induced enhanced permeability to FITC-dextran (A) and reduced TEER (B), respectively, in recipient BMECs. **C**, Representative immunofluorescence staining of ZO-1 in recipient BMECs that, after exposure to *R*-ECExos (*R*-Exo) for 6 hr, were treated with naked miR23a mimic (miR23a), Ctl Oligo-Exos, or miR23a-Exo for 66 hrs. **D**, Exposure to *R*-ECExos in normal media weaken the LBF, indicated by the detachment work (in pJ) between recipient BMECs measured using fluidic AFM at 72 hr post exposure. ***, *p*<0.001. **E**, Compared to Ctl Oligo-Exo, treatment with miR23a ASO-enriched ECExos (ASO-Exo) ameliorate weakened LBFs in *R*-ECExo-treated BMECs, in a dose-dependent manner. *, *p*<0.05; **, *p*<0.01.

#### 4.2

Using immunofluorescence staining of ZO-1, we also demonstrated that post-exposure treatment with miR23a ASO-enriched Exos (1,000 particles/cell) stabilize the paracellular ZO-1 structure from disruption in exposure to *R*-ECExos (**Fig. 5C**).

#### 4.3

The endothelial barrier is maintained by intercellular multi-protein junctional complexes, functionally sealing the lateral space between cells against unbinding forces on the lateral contact sites[56]. Therefore, measuring the lateral binding forces (LBFs) between homogenous cells in response to different stimuli is crucial to understanding the precise biomechanical mechanism underlying intercellular barrier dysfunctions, which is a major outcome of host responses to infections and inflammation. A new biomechanical strategy using AFM-based fluidic force micropipette technology (fluidic AFM) can directly measure live EC-to-EC LBFs in EC monolayers (**Supple Fig. 1**)[57; 58]. We found that the LBFs between living BMECs were reduced after treatment with *R*-ECExos (**Fig. 5D**), providing direct evidence of the biomechanical nature of the endothelial barrier dysfunction during rickettsial infections. We further observed that miR23a ASO-enriched ECExos can stabilize the LBFs between normal recipient BMECs during exposure to *R*-ECExos in a dose-dependent manner (100, 500, and 1,000 particles/cell: **Fig. 5E**).

Collectively, the data presented from these studies suggest that exosomally targeting specific miRs stabilizes endothelial barrier function during rickettsial infection, potentially providing a novel strategy against deadly rickettsioses that affect humans.

## Discussion

In the present study, we first demonstrated that EC-derived Exos have the capacity to deliver oligonucleotide RNAs to normal recipient BMECs in an RNase-abundant environment. Second, we presented evidence suggesting that miR23a in ECExos impairs normal recipient BMEC barrier function, directly targeting TJ protein ZO-1 miRs. To the contrary, information from our in separate studies using a traditional *in vitro* model and a novel single living-cell biomechanical assay, we demonstrated that miR23a ASO-enriched ECExos improved *R*-ECExo-provoked recipient BMEC dysfunction in association with stabilization of ZO-1 in a dose-dependent manner.

Endothelial hyperpermeability is a significant problem in vascular inflammation associated with various diseases, including infections. A key underlying mechanism is increased paracellular leakage of plasma fluid and proteins. Intracellular bacterial infections caused by facultatively intracellular bacteria mycobacteria[59] and listeria[60] and obligately intracellular bacteria rickettsiae[5; 61]can cause dissociation of the paracellular junctions between ECs, leading to transendothelial flux and resulting in microvascular hyperpermeability. Sporn *et al*. reported that bacteria-free conditioned culture media from 24-hour *R. rickettsii-infected* ECs activated normal ECs after an 8-hour exposure, but the results were not statistically significant[62]. We reported that prolonged exposure (72 hrs) to *R*-ECExos induced dysfunction of recipient normal BMECs[26], depending on exosomal RNA cargos and in a dose-dependent manner. *R*-ECExos induced disruption of the TJ protein ZO-1 in recipient ECs[26].

Exo-mediated functional transfer of miRs has been reported with a broad range of downstream effects on recipient cells[45]. EC barrier properties are structurally determined by junctional complexes between ECs, mainly the AJs and TJs[27; 28]. Modulation of EC barrier function by miR23a has been documented by targeting TJs[29; 53]. We found that naked miR23a has little impact on the endothelial barrier function of normal recipient BMECs, likely due to exposure to abundant RNases in normal culture media. However, miR23a-enriched ECExos can markedly induce barrier dysfunction in normal recipient BMECs, suggesting an EV-mediated route for extra-cellular miR23a to play a functional role in the pathogenesis of rickettsial infections, and probably other vascular-targeting infections as well.

Sequence-based ASOs have been shown to have pharmacological effects in the context of infections in a manner dependent upon small noncoding RNAs[54]. However, RNA oligos are quickly digested by RNases both *in vitro* and *in vivo*[63; 64]. Residues of synthetic ASOs are modified to increase their stability, reducing the ability of an oligo to trigger degradation of RNA following hybrid formation[65]. Given the facts that exosomal miR is protected from RNase while free miRNA is degraded immediately and completely by RNases[30; 31; 32], cell-type specific Exos have been developed as novel vehicles to deliver oligos into target recipient cells to exert their functional roles *in vitro* and *in vivo*[30; 50]. Our data suggest that ECExos are an effective vehicle to deliver miR23a ASOs to recipient BMECs, where they neutralize the impact of *R*-ECExos on recipient BMEC barrier function.

Host cell-derived Exos are membrane-derived particles surrounded by a phospholipid bilayer. This inherent property affords Exos with high biocompatibility and reduced toxicity[16]. Exo uptake involves the interplay between Exos and the plasma membrane of the recipient cell[66]. The nature of the endogenous membrane-derived surface enables Exo not only to penetrate the cellular membrane, but also to potentially target specific cell types[67]. Reported evidence suggests that Exos derived from different cells possess different affinities to different cell types. For example, Exos from primary neurons are only taken up by other neurons, whereas those from a neuroblastoma cell line bind equally to astrocytes[68]. Similarly, bone marrow dendritic cell Exos were preferentially captured by splenic dendritic cells, rather than by B or T cells[69]. Exos from oligodendroglia precursor cells were taken up by microglia but not by neurons or astrocytes[70]. By contrast, HeLa cells are able to take up Exos derived from various cells[16]. Furthermore, the surfaces of host cell-derived Exos have an inherently long half-life in circulating blood[71]. In the present study, we employ normal ECExos as the vehicle to deliver ASOs into recipient BMECs. In future we plan to characterize the efficacy of cell-type Exos in blood in the delivery of ASOs because blood has the potential to provide an unlimited source of Exos, although they are heterogeneous in cell type[30; 50].

The biomechanical nature of the barrier between ECs in response to different stimulants is crucial to understanding the precise biomechanical mechanism underlying EC barrier dysfunctions. The epithelial or endothelial barrier is maintained by intercellular multi-protein junctional complexes, either AJs and TJs, functionally sealing the lateral spaces between cells against unbinding forces on the lateral contact sites[56]. The interplay between TJs and AJs regulates major rate-limiting paracellular pathways by allowing particles to permeate across the paracellular route[72] and establish cell polarity[73]. Dysfunctions, ruptures, and breaches of the epithelial or endothelial barriers are major causes of infection and inflammation. Therefore, measuring the LBFs between homogenous cells in response to different stimuli is crucial to understanding the precise biomechanical mechanism underlying intercellular barrier dysfunctions, which is a major outcome of host responses to infections, including inflammation. Traditionally, paracellular permeability can be indirectly evaluated using two methods: measuring TEER[74] and fluorescein tracers after passing through the monolayer[46]; both methods involve indirect measurements[75]. Conventional technologies to directly measure LBFs of living cell-to-cell contacts were not available until a recent report using fluidic AFM for the direct measurement of LBFs on a cell monolayer[76], demonstrating the ability to assess EC LBFs in a mature monolayer in physiological settings, thus providing further evidence that these types of tools can be used to enhance our knowledge of biological processes in developmental biology, tissue regeneration, and disease states like infection and inflammation. During the fluidic AFM assay, we observed that the LBFs between living BMECs was reduced following treatment with *R*-ECExos, providing direct evidence of the biomechanical nature of the endothelial barrier dysfunction during infection with *Rickettsia*. We further observed that miR23a ASO-enriched ECExos can stabilize the LBFs between normal recipient BMECs during exposure to *R*-ECExos in a dose-dependent (100, 500, and 1,000 particles/cell) manner. To our knowledge, this is the first report involving assessment of the mechanical effect of an exosomal RNA cargo on its recipient cell at the level of a living single cell.

Taken together, this study characterizes a functional role for exosomal miR23a in endothelial hyperpermeabilization during rickettsial infection. We found that delivery of miR23a in Exos was efficiently taken up by normal recipient ECs, resulting in impairment to EC barrier structures and functions. Importantly, the observed barrier dysfunction was reversed by the delivery of miR23a ASOs in ECExos to the recipient cells. Future *in vivo* studies will be conducted to explore the potential of Exo-based therapy during bacterial infection and inflammation, and then extend these findings to other infectious pathogens. The ultimate goal is to devise new strategies and to develop novel therapeutics that take advantage of exosome-based biology.

## Materials and Methods

### Nanoparticle tracking analysis

NTA was performed using the TRPS technique and analyzed on a qNano Gold system (Izon, Medford, MA) to determine the size and concentration of EV particles. With the qNano instrument, an electric current between the two fluid chambers is disrupted when a particle passes through a nanopore NP150 with an analysis range of 70 nm to 420 nm, causing a blockade event to be recorded. The magnitude of the event is proportional to the amount of particles traversing the pore, and the blockade rate directly relates to particle concentration that is measured particle by particle. The results can be calibrated using a single-point calibration under the same measurement conditions used for EV particles (stretch, voltage, and pressure)[77]. CPC200 calibration particles (Izon) were diluted in filtered Izon-supplied electrolyte at 1:500 to equilibrate the system prior to measuring EVs and for useused as a reference. Isolated extracellular vesicle samples were diluted at 0, 1:10, 1:100, and 1:1,000. For the measurements, a 35 μl sample was added to the upper fluid chamber and two working pressures, 5 mbar and 10 mbar, were applied under a current of 120 nA. Particle rates between 200 and 1500 particles/min were obtained. The size, density, and distribution of the particles were analyzed using qNano software (Izon).

### Imaging of label-free extracellular vesicles using atomic force microscopy (AFM)

Tapping mode AFM provides a three-dimensional image of surface structures, including height image or deflection image [78; 79; 80], and is commonly used to evaluate the integrity of EVs at the single particle level. To obtain deflection images, the purified and concentrated EV sample was diluted at 1:10, 1:100, and 1:1000 with molecular grade water. Glass coverslip were cleaned three times with ethanol and acetone, then three times with molecular grade water. The coverslip was coated with the diluted EV samples on the designated area for 30 minutes, before being examined using an AFM (CoreAFM, Nanosurf AG, Liestal, Switzerland) using contact mode in the air. A PPP-FMR-50 probe (0.5-9.5N/m, 225μm in length and 28μm in width, Nanosensors) was used. The parameters of the cantilever were calibrated using the default script from the CoreAFM program using the Sader *et al*. method[81]. The cantilever was approached to the sample under the setpoint of 20 nN, and topography scanning was done using the following parameters: 256 points per line, 1.5 seconds per line in a 5-μm x 5-μm image.

### Preparation of oligonucleotide (mimic or ASO)-enriched exosomes

miR23a mimics or ASOs (Integrated DNA Technologies, IDT, oralville, Iowa) were loaded into exosomes using calcium chloride transfection as described[30]. The miRNA mimic or ASO was mixed with exosomes in PBS at 2 pmol oligonucleotides/1 x 10^8^ Exos, followed by the addition of CaCl_2_ (final concentration 0.1 M). The final volume was adjusted to 300 μl using sterile PBS. The mixture was placed on ice for 30 min. Following heat shock at 42°C for 60 secs, the mixture was again placed on ice for 5 min. For the RNase treatment, exosomes were incubated with 5 μg/ml of RNase (EN0531; Thermo Fisher) for 30 min at 37°C as described[30].

### *In vitro* cell model of Exo treatment

Human BMECs (iXCells, SanDiego, California) were cultivated in 5% CO_2_ at 37°C on type I rat-tail collagen-coated round glass coverslips (12 mm diameter, Ted Pella, Redding, CA) or inserts in 24-well plates (0.4 μm polyester membrane, CoStar, Thermo Fisher Scientific) until 90% confluence was achieved[82]. Cells were exposed to *R*-ECExos (1,000 particles/cell) for 6 hrs prior to treatment with exosomes or naked oligonucleotides (Oligo) (IDT) by direct addition of the specified quantity of exosomes or Oligos in the culture media for 66 hrs. Cells were subjected to downstream assays, and all experiments were performed in triplicate.

### Fluidic AFM assay for measuring lateral binding forces (LBF) of BMECs

The LBF is measured using the FluidFM system (Cytosurge, Nanosurf AG, Liestal, Switzerland). A micropipette fluidic AFM cantilever with an 8 μm aperture and spring constant of 2 N/m (CytoSurge) was calibrated using the Sader method in air and liquid by performing deflection and crosstalk compensation. Using an inverted microscope, the cantilever was kept at 20 mbar positive pressure by the fluidic pressure controller (Cytosurge) of the FluidFM system and manipulated to approach the surface of a BMEC in a monolayer. Prior to every measurement, the cantilever was set 45 μm away from the cell. The cantilever approached the target cell at 5 μm/sec until a set point of 20 nN was reached. After a set pause of 10 sec, suction pressure of −200 mbar decreased to −400 mbar to ensure a seal between the apical surface of the cell and the aperture of the hollow cantilever. The cantilever was vertically retracted to a distance of 100 μm to separate the targeted BMEC from the substrate and the monolayer, and the unbinding effort was assessed by measuring the work done (in picojoules [pJ]), which was calculated by integrating the area under the force-distance (F-D) curve using software as we described[83; 84]. The captured BMEC was released by applying 1,000 mbar positive pressure prior to moving the cantilever to another individual BMEC that did not contact other cells in the same culture. The LBF value was obtained using the method of Sancho *et al*. [76] by measuring the unbinding force/work required to separate the individual BMEC from the substrate. The Temperature Controller (NanoSurf) kept the liquid environment at 37°C during all measurements.

### Statistical analysis

All data were presented as mean± standard error of the mean and analyzed using SPSS version 22.0 (IBM^®^ SPSS Statistics, Armonk, NY). A two-tailed Student’s *t*-test or one-way analysis of variance (ANOVA) was used to explore statistical differences. If the ANOVA revealed a significant difference, a post hoc Tukey’s test was further adopted to assess the pairwise comparison between groups. The level of statistical significance for all analyses was set at *p*<0.05.

## Acknowledgements

We gratefully acknowledge Dr. Kimberly Schuenke for her critical review and editing of the manuscript. We thank Drs. Bao X and Fang R for reagent supports during the planning phases of the experiments. This work was supported by NIH grant R01AI121012 (BG), R21AI137785 (BG), R21AI154211(BG), R03AI142406 (BG), R21AI144328 (BG) and John Sealy Distinguished Chair in Alzheimer’s diseases (XF). The sponsors had no role in the study design, data collection and analysis, decision to publish, or preparation of the manuscript.

## Authorship Contributions

CZ, JB, and YQ performed experiments, analyzed data, and prepared manuscript. QC and EN performed experiments and analyzed data. JY, BK, KK, and YJ designed the study. XF and AG designed the study, analyzed data, and prepared the manuscript. BG designed the study, performed experiments, analyzed data, and wrote the manuscript.

